# The diminished ovarian reserve with chronic stress induced by tripterygium with chronic unpredictable mild stress in SD rats

**DOI:** 10.1101/2023.05.12.540547

**Authors:** Yanhua Li, Wei Wang, Jun Liu

## Abstract

**Aim:** To establish a rat model of DOR combined with chronic unpredictable mild stress (CUMS), and to explore the effect of chronic adverse psychological stress on ovarian reserve dysfunction, so as to provide experimental basis for further research.

**Methods:** The rats were randomly divided into control group, CUMS group, DOR group and DOR +CUMS group, with 10 rats in each group,given fed normally, chronic mild unpredictable stress, Tripterygium glycoside tablets and Tripterygium glycoside tablets with CUMS intervention in proper order for 21 days. We observed the body mass, estrous cycle, behavioral testing, the hematoxylin and eosin (H&E) staining of ovarian tissue,the serum levels (E2, FSH, 5-HT and GnRH)and Apoptotic granulosa cells of ovarian tissue.

**Results:** After an 21-days exposure, in DOR group or DOR+CUMS group the serum levels of E2 was reduced and the FSH was raised. In DOR group, the estrous cycle was disordered, the ovary was slightly atrophied and the ovarian tissue structure was not clear, the number of follicles and the content of follicular fluid were small, the cell layers was reduced, and inflammatory cell infiltration was visible, the change was more obvious in DOR+CUMS group. In CUMS group and DOR+CUMS group, the OFT, SPT and 5-HT were lower and the GnRH was higher than in control group. About the TUNEL, the AI in the DOR group, the CUMS group and DOR+CUMS group were higher.

**Conclusions:** Our outcomes state that we could established an rat model of psychological stress-induced DOR successfully which can be used for further study.

## 1. INTRODUCTION

Diminished ovarian reserve (DOR) refers to the phenomenon of decreased fertility due to both of the quantitative and qualitative reduction in oocytes. DOR can lead to the decreased menstruation, sparse menstruation, or premature ovarian failure [1]. DOR is a female endocrine disorder with many pathogenic factors including heredity, environment, immunity, psychology, living habit, etc[2]. A large amount of evidence has shown that DOR patients have a high rate of psychological disorders such as anxiety and depression, and adverse psychological stress[3]. Furthermore, diseases usually interact to each other to form a bad cycle. Therefore, the effect of chronic psychological stress on DOR has attracted widespread attention[4-5]. However, due to the lack of a proper animal model which bears the similar symptoms of DOR, it is hard to know the effect mechanism clearly.

Chronic unpredictable mild stress (CUMS) is widely used in inducing anxious-like behavior in animal models and researching chronic-stress-associated illness. CUMS mainly includes stressful stimuli which mimics the stress of daily life. This model has been employed to the study of the neurobiological processes that adjust the influence of chronic stress [6]. However, the studies on ovarian after stress exposure through the kind of model have been hardly reported[7]. As well as, few of studies have induced DOR phenotype and explored the serum hormone changes being related to DOR.

Hence, it is particularly important to build a proper animal model and reveal the internal mechanism of neuroendocrine and reproductive endocrine in the DOR pathogenesis[8]. In this work, the rat model of DOR was prepared by Tripterygium wilfordii polyglycoside tablets. On this basis, the rat model of DOR combined with chronic stress was established by CUMS. Furthermore, the model was analyzed and evaluated according to the reproductive hormone changes and follicle development-related factors. And the chronic stress damage on the ovarian biological functions was further studied.

## 2. MATERIALS AND METHODS

### 2.1 Animals

Forty specific-pathogen-free (SPF), female Sprague–Dawley (SD) rats (aged 12 weeks, weighing 200±10g) exhibiting regular 4–5-day estrous cycles were obtained from Beijing Vital River Laboratory Animal Technology (animal license: SCXK (jing) 2016-0006). The rats were provided free access to food and water and were group-housed in the animal laboratory of Beijing University of Chinese Medicine at the temperature of 20–25°C and the relative humidity of 40–60% on a 12 h/12 h light/dark cycle. The experimental procedures were performed in strict accordance with the recommendations of the Guide for the Care and Use of Laboratory Animals issued by the U.S. National Institutes of Health, and the study was approved by the Ethics Committee of Beijing University of Chinese Medicine (No. BUCM-4-2019030408-1125). Two normal consecutive estrous cycles were used in the work, following a week of adaptation to the laboratory conditions. The rats were divided into the following four groups: a control group (n=10), a CUMS group (n=10), a DOR group (n=10), and a DOR combined with CUMS group (n=10).

### 2.2 Grouping and model establishment

The rats were randomly divided into a Control group, a CUMS group, a DOR group and a CUMS combined with DOR (DOR+CUMS) group. There were 10 rats in each group. Rats in the Control group were housed together (six rats per cage), received water and food ad libitum, and were not exposed to the CUMS paradigm. The DOR model group was administered Tripterygium wilfordii polyglycoside tablets, at a concentration of 50 mg/kg/d[9]. Tripterygium wilfordii polyglycoside powder was dissolved in 1% carboxymethylcellulose (CMC)-Na and diluted to a final Tripterygium solution concentration of 6.25 mg/ml. The rats were administered a volume of 8 mL/kg of this solution via gavage once daily for 21 days. The CUMS paradigm involved the use of ten different stressors[10], which were randomly applied each week over a total of 3 weeks. The stressors were as follows: (1) solitary cage feeding; (2) fasting for 24 h; (3) water deprivation for 24 h; (4) tail clamping for 1 min; (5) Black and white upside down (12 h dark, 12 h daytime); (6) exposure in a wet cage for 24 h; (7) swimming in cold water (4°C) for 5 min; (8) 45° cage tilting lasting for 24 h; (9) foreign body stimulation for 24 h (such as hardwood stick, rag, etc.); (10) restricted movement for 3 h. The modeling method of DOR+CUMS group was the same as DOR and CUMS groups.

### 2.3 General situations

The changes of defecation, activity, hair color, spirit, water intake and other general conditions of rats every day was observed and recorded. The body mass once a week was weighed and the change of body mass before the experiment (after the end of adaptive feeding), on the 7th, 14th and 21st day was compared in each group.

### 2.4 Estrous cycle

After the beginning of the experiment, the exfoliated cells in the vagina of rats were smeared at 10 a.m. every day. 30 μl of normal saline was sucked with a pipette and injected into the vagina of rats, which was repeated 3-5 times. The sucked liquid was applied on the pre-marked slide. After drying, the exfoliated cells on the slide were fixed with 95% ethanol for 30 min. Then, the exfoliated cells were stained with the Wright Giemsa compound dye and observed by the optical microscope. The estrous cycle of rats including normal, prolonged, stagnant and no cycle was determined by the continuous smears[11].

### 2.5 Behavioral tests

The behavioral tests were performed by three trained and experienced observers who were blinded to the treatment conditions[12-13]. All animals were subjected to the behavioral tests after 21 days of drug or distilled water administered via gavage.

### 2.5.1 Open field test (OFT)

The inner surface of the OFT apparatus was painted black. The floor of the apparatus (53×59×48 cm^3^) was divided into a grid of 25 identical squares (20×20 cm^2^) with white stripes. A rat was placed at the center of the arena and freely allowed to explore for 3 min in a dark and quiet room. Three or four paws crossing the line, or more than half of the body of the rat entering an adjacent square was defined as crossing the square. The number of squares crossed by the rat was counted as the horizontal score. The frequency at which two forelimbs of the rat left the ground was counted as the vertical score. The total OFT score was calculated as the sum of the horizontal and vertical score. After each test, the apparatus was fully cleaned with 75% ethanol.

### 2.5.2 Sucrose preference test (SPT)

All animals were food-deprived for 24 h before testing. Subsequently, two bottles containing either 1% sucrose solution (w/v, sucrose dissolved in distilled water) or distilled water were put in the cage for the rats to consume on the test day. After 1 h, the volume of consumption of each solution was recorded. And the data was analyzed based on the formula: Percentage sucrose preference= (volume of sucrose solution consumption/total fluid intake volume) × 100%.

### 2.6 Specimen collection and processing

The rats of each group were fasted with water and food for 12 h after OFT and SPT. Then the rats were weighed and anesthetized with the 50 mg/kg dose of 1% Pentobarbital Sodium. The blood extracted from the abdominal aorta was kept for 2 h at the room temperature. Then the blood was centrifuged at 3000r/min for 10 min. Subsequently, the supernatant was derived and stored in the refrigerator with -20 °C. The bilateral ovaries were taken out and the attached other tissues were also removed. Then the bilateral ovaries were weighed, recorded, and stored in the liquid nitrogen with -80 °C.

### 2.7 Enzyme linked immunosorbent assays (ELISA)

The concentration of follicle-stimulating hormone (FSH), Estradiol (E2), 5-hydroxytryptamine (5-HT) and gonadotropin-releasing hormone (GnRH) was determined using ELISA kits (96T, TSZ, USA) on the basis of the manufacturer’s instructions. The brief processes included samples adding, incubating, washing, adding diluted biotinylated anti-immunoglobulin G (IgG), incubating, washing, HRP and measuring OD[14].

### 2.8 Histological observation of ovary

The ovarian morphology and tissue structure was acquired by the optical microscope[15]. The number of follicles at all levels was counted. The experimental procedure was as following. Firstly, the slices were baked at 60 °C for 15 min. Secondly, the slices were dewaxed with xylene for two consecutive times which were 15 min each time. Thirdly, the slices were hydrated by the gradient alcohol (ethanol concentration: 100%-90%-75%-50%, 10 min each time), and soaked in distilled water for 5 min. Then the slices were soaked by the 0.01 mpbs (pH=7.4) for 3 times which were 5 min each time. Subsequently, the slices were seriatim stained with hematoxylin, incubated at room temperature for 10 min, differentiated for 2-3 s with 1% hydrochloric acid alcohol, rinsed with tap water to return to blue, soaked with 0.01 mpbs (pH=7.4) for 3 times (5min each time), stained with eosin at room temperature for 3 s, washed with absolute alcohol, soaked with xylene twice (10 min each time), sealed with neutral gum, and observed by the microscope.

### 2.9 Terminal deoxyribonucleotidyl transferase-mediated dUTP–digoxigenin nick end-labelling (TUNEL)

The TUNEL staining was conducted as following. The ovarian sections with embedded 4 μm paraffin were deparaffinized with xylene, rehydrated with ethanol, incubated with 20 μg/mL proteinase K at 37 °C for 20 min, incubated with recombinant TdT reaction mixture at 37 °C for 60 min in a humidified dark chamber, and washed with PBS. Then, the Converter-POD was added into the dealt ovarian sections[16]. Subsequently, the ovarian sections were incubated for 30 min at 37 °C, dehydrated, fixed on slides. The brown nuclear stain represents apoptotic cells under the microscope. The percentage of apoptosis granulosa cells was observed from the random high-power fields (×200) of each slide to determine the apoptotic index (AI).

### 2.10 Statistical analysis

The experimental data were processed and analyzed by Statistical Package for the Social Sciences (SPSS, IBM Corporation, NY, USA) version 23.0. The measured data are shown as the means±standard deviations (SD). The data which do not follow the normal distribution were compared by nonparametric test. The data obeying the normal distribution were compared by one-way analysis of variance(ANOVA) followed by two-sided Student’s test. While the variance is homogeneous, the LSD method was employed to analyze. Or else, the Dunnett’s method was used. The differences of *p*<0.05 were considered statistically significant.

## 3. RESULTS

### 3.1 Effects of intervention of Tripterygium combined with CUMS on the body mass in model rats

Figure 1 exhibits the dependent relations between the body mass and days in the model rats. Obviously, there was no significant difference in body mass of each group before and after 7 days modeling (*P*>0.05). After 14 days of modeling, the body mass in CUMS group was smaller than that of the Control group, and that the DOR+CUMS group was lower than that of the CUMS group (*P*<0.05). After 21 days of modeling, the body mass in the DOR+CUMS group was the lowest among the all groups (*P*<0.05). Furthermore, the increment of the body mass of the DOR+CUMS group was much smaller than that of the Control and CUMS groups. As a result, the intervention of Tripterygium combined with CUMS reduced the body mass in model rats.

**Figure 1.**
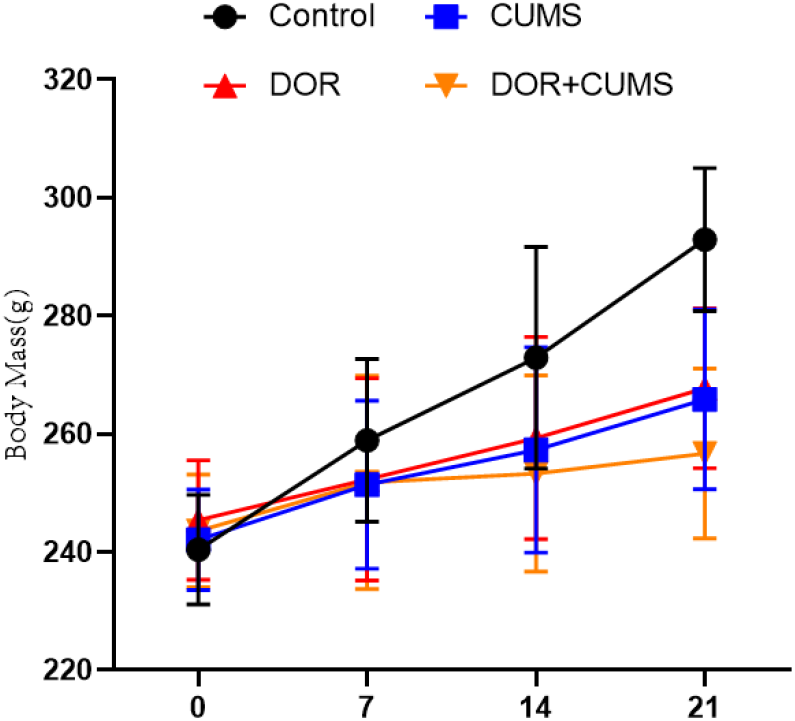
Day dependence of body mass in model rats.

### 3.2 Effects of intervention of Tripterygium combined with CUMS on the estrous cycle disordered in model rats

The estrous cycle of model rats was monitored every day. All model rats were normal before the experiment. Figure 2 shows the disordered estrous cycle for the Control, CUMS, DOR, and DOR+CUMS groups after 21 days modeling. It can be observed that the estrous cycles of model rats in DOR and DOR+CUMS group were disordered (*P*<0.05), where the estrous cycle was prolongation, stagnation, and no cycle. However, the Control and CUMS groups shown no significant difference in the estrous cycle (*P*> 0.05). So, the **intervention of Tripterygium** combined **with CUMS** promoted **the estrous cycle disordered in model rats**.

**Figure 2.**
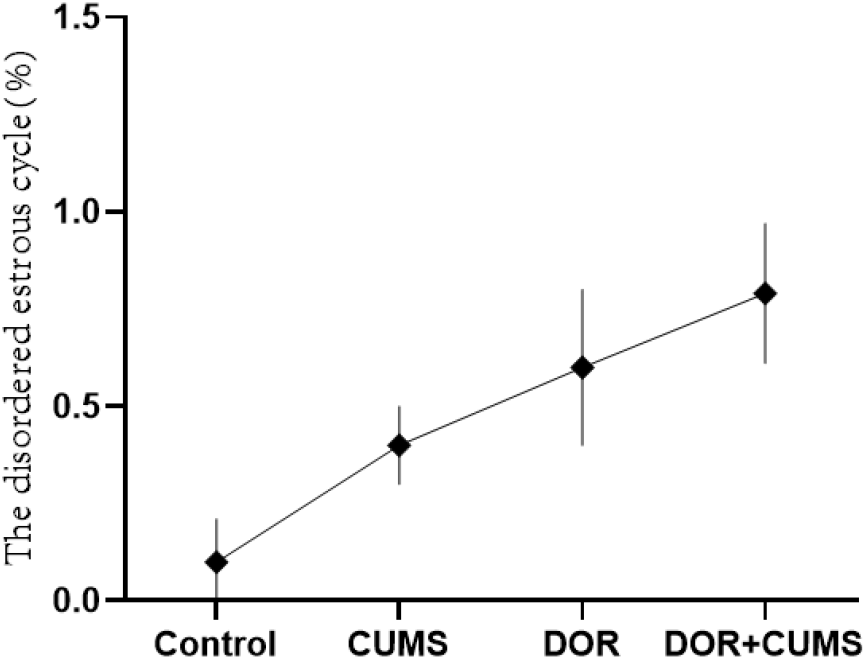
The disordered estrous cycle for the Control, CUMS, DOR, and DOR+CUMS groups.

### 3.3 Effects of intervention of Tripterygium combined with CUMS on the psychological behavior in model rats

The chronic psychological stress-like behaviors were evaluated via the OFT and SPT behavioral tests. As shown in Figure 3A, the CUMS and DOR+CUMS groups presented the much lower score of the central total distance than that Control group on the 21th day (*P*<0.05). In addition, the DOR group exhibited no significant differences (*P*>0.05). In comparison to the CUMS group, the score of the central total distance for the DOR+CUMS group was no obvious differences (*P*>0.05). The SPT results of four group of model rats were displayed in Figure 3B. Conspicuously, the CUMS and DOR+CUMS groups exhibited much lower sucrose preference than that Control group on the 21th day (*P*<0.05). And the DOR group shown no distinct differences (*P*>0.05). Additionally, in comparison to the CUMS group, the SPT scores of the DOR+CUMS group were no prominent differences (*P*>0.05). As a result, the intervention of Tripterygium combined with CUMS leaded to the chronic psychological stress-like behavior in model rats.

**Figure 3.**
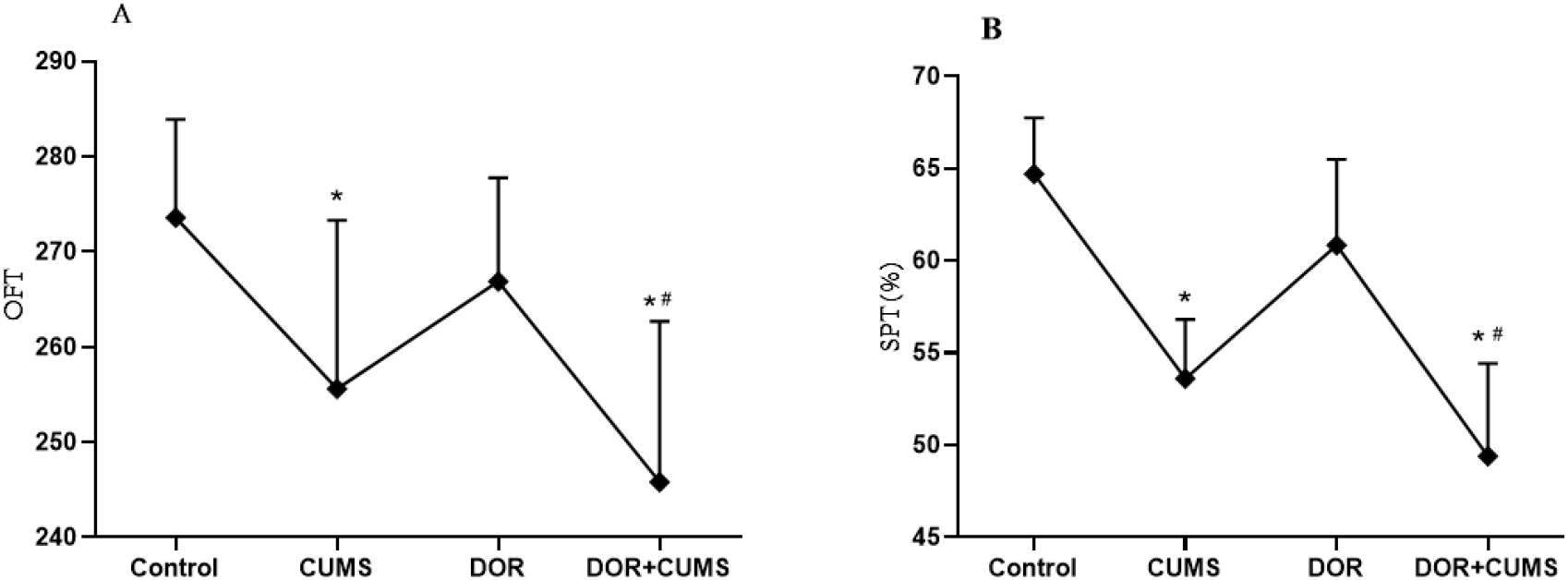
The influence of Tripterygium combined with CUMS on the psychological behavior of rats (N=10/group). (A) The central total distance scores in the open field test (OFT) and (B) sucrose preference test (SPT) results of the Control, CUMS, DOR, and DOR+CUMS groups after modeling. The data were presented as the mean ± SD. ^*^*p*<0.05 vs. control group; # *p*<0.05 vs. DOR group.

### 3.4 Effects of intervention of Tripterygium combined with CUMS on the serum hormone levels in the model rats

The serum hormones were measured by ELISA (N=10/group). Figure 4 shown the serum levels of FSH, E2, 5-HT, and GnRH. From Figures 4A and 4B, the FSH of DOR and DOR+CUMS groups was higher than that of Control and CUMS groups (*p*<0.05). In contrast, the F2 of DOR and DOR+CUMS groups was lower (*p*<0.05). Furthermore, there was no obvious difference between DOR and DOR+CUMS groups in FSH and E2 (*p* > 0.05). From Figures 4C and 4D, the CUMS and DOR+CUMS groups delivered smaller 5-HT than that Control and DOR groups (*p*<0.05). On the contrary, the GnRH in CUMS and DOR+CUMS groups was bigger (*p*<0.05). However, there was no significant difference between CUMS and DOR+CUMS groups in GnRH and 5-HT (*p* > 0.05). As a result, the intervention of Tripterygium combined with CUMS disrupted the serum hormone levels of the model rats.

**Figure 4.**
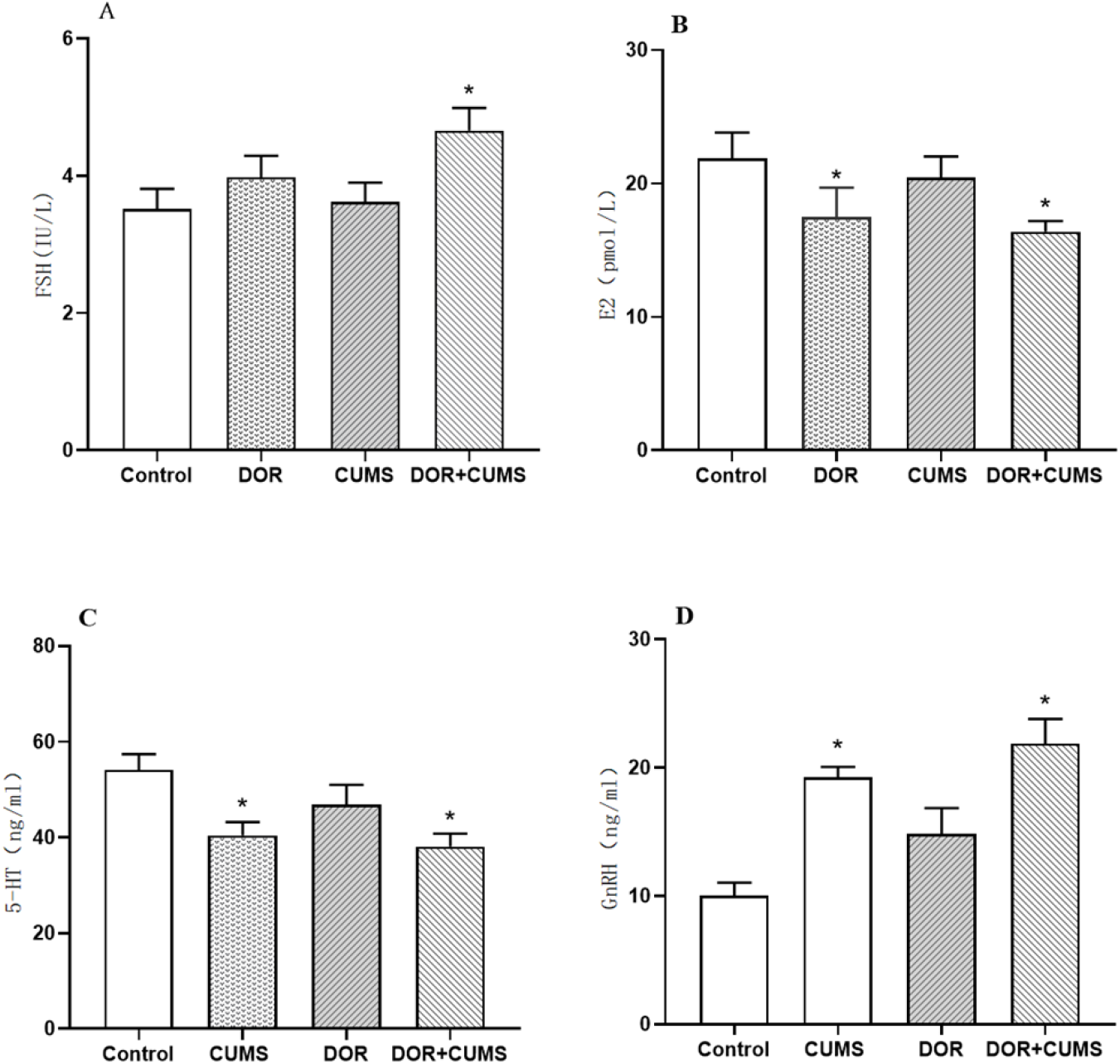
Serum levels of (A) follicle-stimulating hormone (FSH), (B) estradiol (E2), (C) 5-hydroxytryptamine(5-HT), and (D) gonadotropin-releasing hormone (GnRH). The data are presented as the mean±SD. ^*^*p*<0.05 vs. control group.

### 3.5 Effects of intervention of Tripterygium combined with CUMS on the development of follicles in the model rats

Figure 5 shown the changes in the pathophysiology of ovarian tissue for the the Control, CUMS, DOR, and DOR+CUMS groups. The hematoxylin & eosin was employed to stain. In Figure 5A, not only the ovarian tissue structure was clear, but also the morphology of follicles at all levels was complete in the Control group. And that the blood vessels, nerves, and lymphatic vessels were plentiful in the Control group. There were also no abnormalities of fibrosis and interstitial inflammation. Furthermore, we could observe more primordial follicles, primary follicles, secondary follicles, mature follicles, and fewer atretic follicles in Figure 5A. In comparison to the Control group, the CUMS group exhibited fewer normal follicles and corpus luteum. According to Figures 5C and 5D, the DOR and DOR+CUMS groups revealed cortical hyperplasia, medullary atrophy, interstitial fibrosis, untight and out-of-order arrangement of granulosa cells, and decreased quantity of all levels follicles. Consequently, the intervention of Tripterygium combined with CUMS restrained the development of follicles in the model rats.

**Figure 5.**
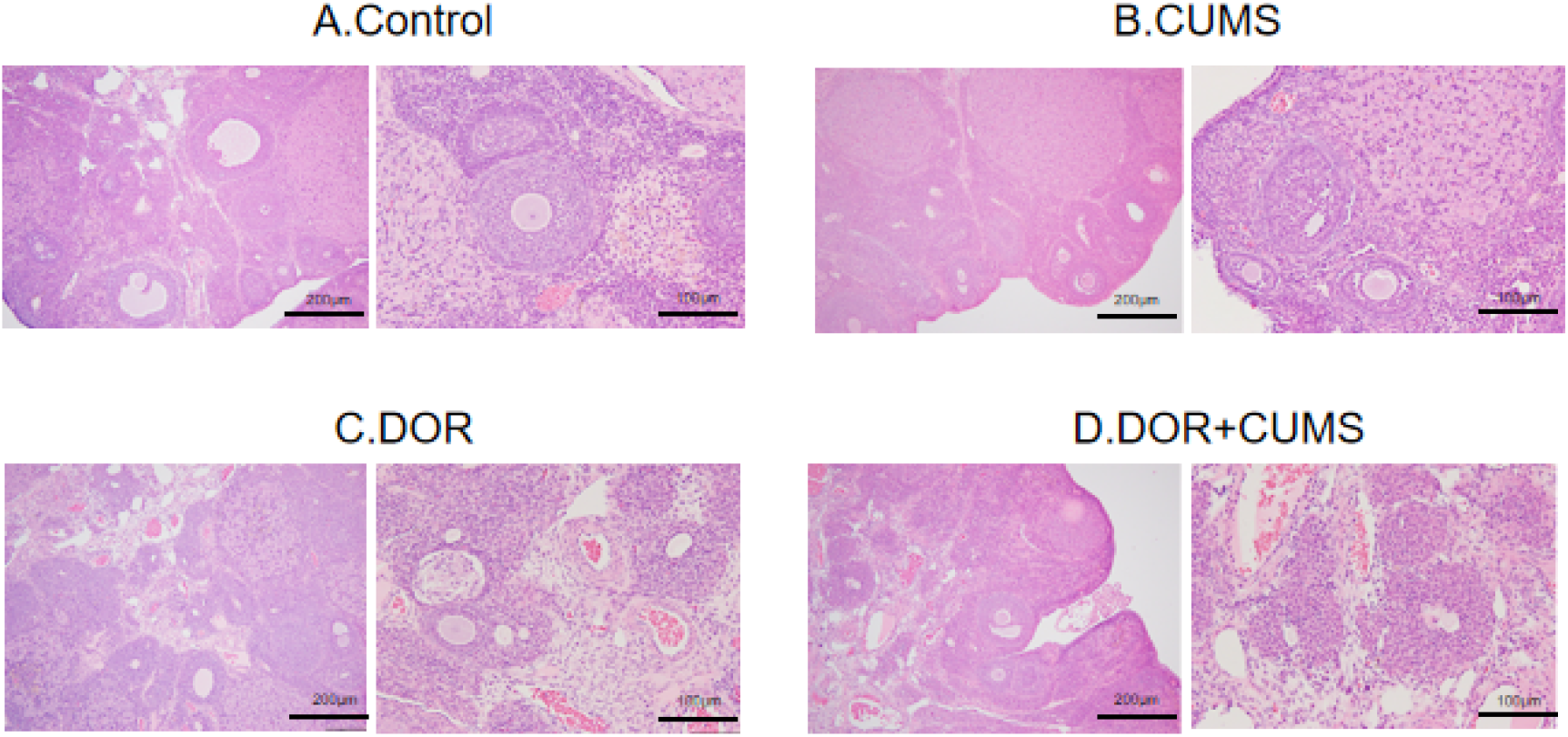
Changes in the pathophysiology of ovarian tissue (hematoxylin & eosin staining, magnification=200×, 100×). The Optical micrographs show the ovarian tissue from the Control (A), CUMS group (B), DOR group (C), and DOR+CUMS groups (D) (N=10/group).

### 3.6 Effects of intervention of Tripterygium combined with CUMS on apoptosis of ovarian granulosa cells in the model rats

The TUNEL staining was used to estimate the apoptosis of ovarian granulosa cells. Figure 6A exhibits the optical microscope images of ovarian tissue sections stained by terminal deoxynucleotidyl transferase dUTP nick end labeling (TUNEL). The red images represented the nuclei of TUNEL-positive cells in Figure 6A. According to Figure 6B, the apoptotic index of DOR and DOR+CUMS groups was bigger than that of the Control group (*p*<0.05). Additionally, the apoptotic index of the CUMS group was slightly higher than that of Control group, but there was no statistically conspicuous difference (*p* > 0.05). As a result, the intervention of Tripterygium combined with CUMS accelerated apoptosis of the ovarian granulosa cells.

**Fig.6.**
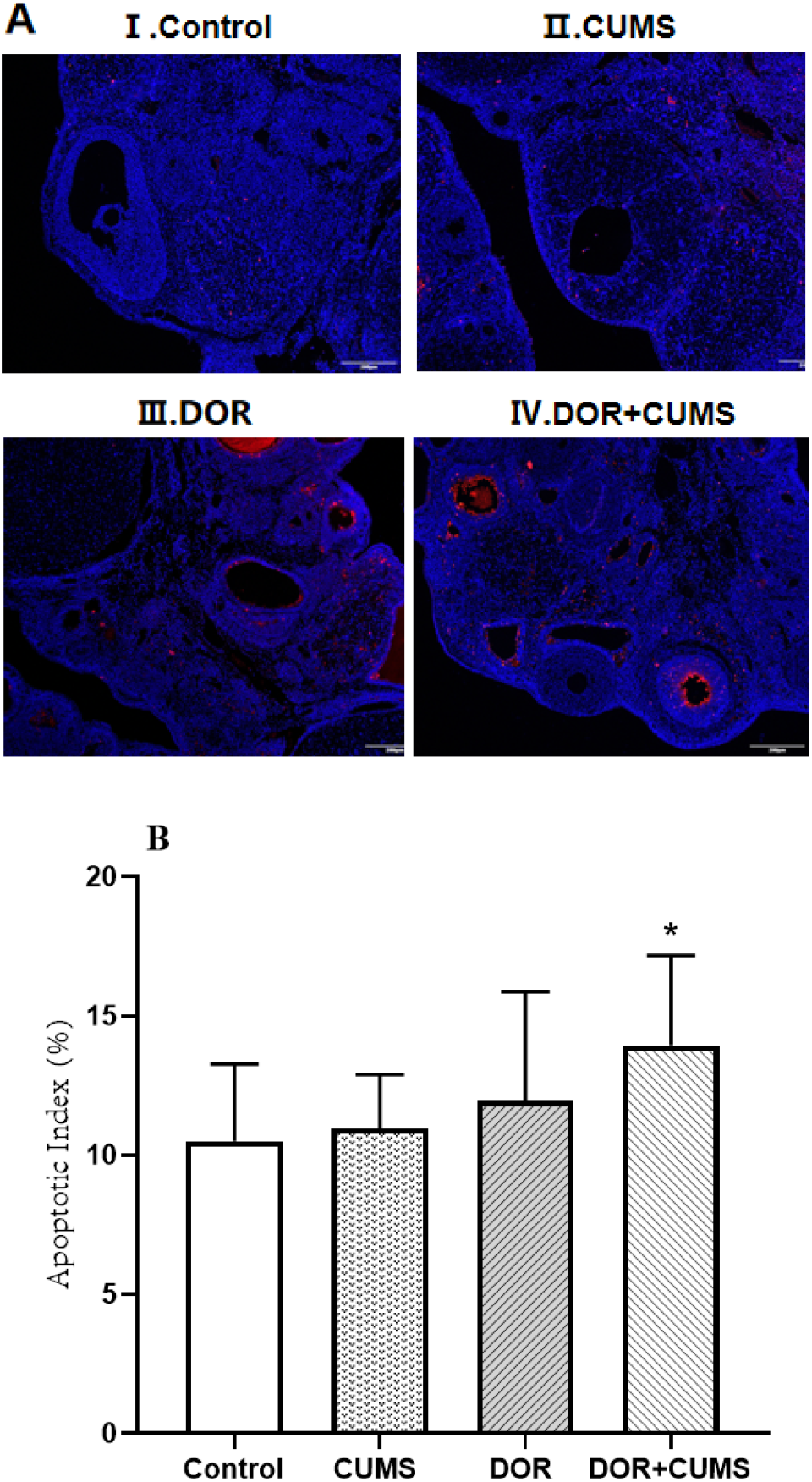
The influence of Tripterygium combined with CUMS on apoptosis of ovarian granulosa cells of rats. (A) Optical images of ovarian tissue sections stained with terminal deoxynucleotidyl transferase dUTP nick end labeling (TUNEL): I. Control group, II. CUMS group, III. DOR group, IV. DOR+CUMS group. (magnification= 200×, scale bars= 200 μm). (B) The apoptotic index of each group (N=10/group). The data are presented as the mean±SD. *p<0.05 vs. the normal control group.

## 4. DISCUSSION

Up to now, the reasons of DOR is still unclear. Previous studies reported that the DOR was related to heredity environmental factors such as genes and hormones in the body, region, nutrition and lifestyle element correlation [17]. Furthermore, the more and more bad mental state was also reported. It has been demonstrated that the DOR accelerate the occurrence and development of the disease. Therefore, the DOR is not only a reproductive endocrine, and metabolic disorder, but also a physical and mental disease. So, the DOR should be sufficiently concerned in the clinical practice. Consequently, it is quite crucial to establish more animal models close to clinic for exploring pathogenesis and treatment [18].

CUMS was usually employed to construct an animal model of chronic stress [19]. The Tripterygium Wilfordii preparation as a kind of immunosuppressant was extensively used in the clinic. The tripterygium wilfordii preparation delivers reproductive toxicity and leads to varying degrees of damage to ovarian function [20]. Tripterygium Wilfordii affected the reproductive system of female animals during sexual maturity, for examples, the decline of ovarian function, premature ovarian failure, and estrous cycle turbulence [21]. And the influencing mechanism refers to anti apoptotic proteins (Bcl-2, Akt), signal transduction related genes Wnt4, Pro apoptotic proteins (Caspase-3, Bax), inhibiting follicular growth, and inducing ovarian cell apoptosis [22].

The important characteristic changes of DOR were the lessening of the quantity of follicle and reproductive endocrine dysfunction. And the important features of liver depression were the variation of neurotransmitters, OFT, and SPT. For the fundamental study of the DOR, the local histopathological changes of ovary, the levels of serum sex hormones, neurotransmitters and behavioral changes were defined as the observed indicator. In this work, the results shown that the DOR group of model rats exhibited the decreased serum levels of E2, increased FHS levels, out-of-order estrous cycle, slightly atrophied ovary, unclear ovarian tissue structure, low number of follicles, small content of follicular fluid, lessened number of granular cell layers, deformation of cumulus. Furthermore, a little of interstitial blood vessels were dilated, and inflammatory cell infiltration was eyeable in the DOR group. Particularly, those changes of ovarian condition in the DOR+CUMS group were more conspicuous. The CUMS and DOR+CUMS groups delivered much lower OFT, SPT, and 5-HT, and higher GnRH than that Control group. In addition, both of the TUNEL and apoptotic index of the CUMS and DOR+CUMS groups was higher than that of the DOR group.

So far, there are few reports on the composite model of ovarian reserve dysfunction and chronic stress [23]. The preliminary study of this composite model was realized in this work, but there are still some challenges remaining in this area. For example, a future research lies in the interaction between the two modeling methods, and the effective indicators to verify the composite model to provide some ideas for the clinical treatment of female ovarian reserve function decline.

## 5. CONCLUSION

In summary, the animal model was established through the Tripterygium wilfordii polyglycoside tablets combined with CUMS. According to the behavioral changes and neuroendocrine level of rats, the animal model was consistent with the pathological and biochemical characteristics of DOR in terms of the ovarian histomorphology and sex hormone level. Furthermore, the animal model also conformed to the characteristics of stress state, indicating the preliminary construction of the DOR with chronic stress state in this study. This work not only provides a new idea for the research of DOR with adverse psychological stress, but also is of great significance to the pathogenesis of DOR and pharmacodynamic evaluation.

